# Lipid transfer by ORP3 is required for the regulation of PI4P and PI(4,5)P2 at the plasma membrane in mitosis

**DOI:** 10.1101/2025.10.22.684039

**Authors:** Anaïs Vertueux, Agathe Verraes, Christie Ouaddi, Hugo Siegfried, Emilie Pellier, Véronique Proux-Gillardeaux, Laurence Walch, Mélina Heuzé, Cathy L. Jackson, Jean-Marc Verbavatz

**Author notes:** deceased (Véronique Proux-Gillardeaux).

## Abstract

During mitosis, cellular contents including the genetic material and membrane-bound organelles must be faithfully distributed between the two daughter cells. Regulation of PI(4,5)P2 levels at the plasma membrane is essential for mitotic progression, including anchoring of the mitotic spindle, recruitment of the actomyosin cytoskeleton at the cleavage furrow, and abscission. Here, we demonstrate that the ORP3 lipid transfer protein, which transfers PI4P from the plasma membrane to the endoplasmic reticulum (ER) at ER-plasma membrane contacts, plays a crucial role in the regulation of PI4P and PI(4,5)P2 levels at the plasma membrane in mitosis. We show that defects in ORP3 function alter PI4P and PI(4,5)P2 distributions, distribution of the actin cytoskeleton at the plasma membrane, mitotic spindle geometry, chromosome segregation, abscission, and lead to the accumulation of multinucleated cells. The function of ORP3 in mitosis is dependent on its ER-partner VAPA and phosphorylation of the ORP3 VAPA-binding motif strongly recruits ORP3 to the ER, priming it for PI4P transfer from the plasma membrane to the ER. Finally ORP3 is required to prevent PI4P accumulation at the cytoplasmic bridge as a result of PI(4,5)P2 hydrolysis for abscission and successful completion of cell division. Altogether, ORP3 plays a key role in PI4P and PI(4,5)P2 regulation during mitosis. Impairment of ORP3 function results in multiple cell division phenotypes, leading to genetic instability and aneuploidy.

## INTRODUCTION

In eukaryotes, the identity of cell membranes is defined by their specific lipid and protein composition (Jackson et al., 2016). Although phosphoinositides represent a minor fraction of membrane lipids (less than 1% of total phospholipids, Cauvin and Echard, 2015), they play important roles in cellular function and their distribution accross membranes is exquisitely regulated. Phosphoinositides are phosphorylated derivatives of phosphatidylinositol (PI), an abundant membrane lipid. They are formed by the action of specific kinases and phosphatases at position 3, 4 or 5 of the inositol ring (Mandal 2020). The resulting phosphoinositide diversity (7 species) and polarization at membranes constitutes the basis of cell membrane identity, a determinant of cellular functions such as trafficking, cytoskeleton organization, signaling and polarity (Posor et al., 2022; Bugda Gwilt and Thiagarajah, 2022). Phosphoinositide patterns in mitosis are crucial to regulate and anchor the actomyosin and microtubule cytoskeleton at the plasma membrane through adaptor proteins (Cauvin and Echard, 2015). PI(4,5)P2 is the most abundant phosphoinositide at the plasma membrane where it binds a plethora of intracellular proteins (Mandal 2020), including actin-binding and microtubule-associated proteins (Senju and Lappalainen 2019; Gervais et al., 2008; Kotak et al., 2014). During mitosis, starting from prometaphase, thickening of the actin cortex is required to increase cell stiffness in order to support the forces required for cell elongation, chromosome segregation and eventually cytokinesis and abscission. ERM proteins (Ezrin, Radixin, Moesin) bind PI(4,5)P2 at the plasma membrane and are also activated by PI(4,5)P2 (Solinet et al., 2013). This is essential to anchor the cortical actin cytoskeleton, particularly in mitosis (Fehon et al., 2010). This is a highly regulated mechanism, as ERM proteins hyperactivation impairs cell elongation in anaphase due to increased cortex stiffness (Roubinet et al., 2011), whereas ERM inhibition prevents cell rounding and impairs spindle assembly and chromosome segregation (Kunda et al., 2008). ERM activation in early mitosis is necessary for the localization of the LGN-NuMA complex which defines the orientation of the spindle (Machicoane et al., 2014). In addition to ERMs, PI(4,5)P2 can directly recruit NuMA to the cortex at the cell poles where PI(4,5)P2 is concentrated (Kotak et al., 2014). During cytokinesis, PI(4,5)P2 accumulates at the ingression furrow, where it recruits components of the contractile ring. Prior to abscission, ERM proteins are inactivated and PI(4,5)P2 is hydrolyzed into PI4P at the site of abscission in order to depolymerize the actin cytoskeleton and allow for the recruitment of the ESCRT complex (Cauvin and Echard, 2015). Altogether, PI(4,5)P2 distribution at the plasma membrane is tightly regulated in space and time, and determines mitotic progression from prometaphase to abscission.

PI(4,5)P2 mainly results from the phosphorylation of PI4P by PI4P 5-kinases (PI4P-5K), and to a lesser extent from the hydrolysis of PI(3,4,5)P3. At the plasma membrane, PI4P largely originates from vesicle trafficking in the secretory pathway. But PI4P can also result from the phosphorylation of phosphatidylinositol by PI-4 kinases (PI-4K). Another source of membrane phosphoinositides comes from non-vesicular transfer of lipids by lipid transfer proteins. Lipid transfer proteins are recruited at sites of contact between the membranes of distinct organelles, where they transfer lipids from one organelle membrane to the other, using a hydrophobic pocket (Weber-Boyvat et al., 2015; Reinisch and Prinz 2021). These membrane contacts often involve the endoplasmic reticulum (ER), which is the primary organelle of lipid synthesis (Yang et al., 2018).

ORPs (OSBP-related proteins) are one of the largest and best-studied families of lipid transfer proteins. The family has 12 members in humans (Delfosse et al., 2020). In mammals, most ORP proteins harbor 3 important domains (see also Fig. 2A): i) a FFAT motif (two phenylalanines “FF” in an acidic tract), which specify ORP interaction with the MSP domain of VAP membrane proteins at the ER; ii) a “Plextrin Homology” (PH) domain, which binds membrane phosphoinositides and sometimes proteins on the target membrane; and iii) a hydrophobic lipid pocket (ORD domain) for the transfer of a lipid from one membrane to the other (Olkkonen, 2015; Delfosse et al., 2020). PI4P is a conserved substrate among ORP proteins (Jackson et al., 2016, Delfosse et al., 2020). PI4P transport by ORP proteins from the trans-Golgi, endosomes or the plasma membrane where it is abundant, to the ER where its concentrations are low, serves as the driving force for the counter-transport of another lipid (cholesterol, phosphatidylserine) against its gradient (Mesmin et al., 2013; Moser von Filseck et al., 2014; Chung et al., 2015; Mesmin et al., 2017; Antonny 2018). Sohn et al. (2018) have proposed that the counter transport of PI4P and phosphatidylserine by ORP5 or ORP8 at ER-plasma membrane contacts could down regulate PI4P at the plasma membrane. Through the inter-conversion between PI4P and PI(4,5)P2 by kinases and phosphatases, PI4P transport to the ER by ORP5/ORP8 was proposed to regulate plasma membrane PI(4,5)P2 levels as well. However, this mechanism of phosphoinositide regulation at the plasma membrane is dependent on the availability of the counter-transported lipid at the ER.

Interestingly, one member of the ORP family, ORP3, localizes to ER-plasma membrane contacts and transfers PI4P from the plasma membrane to the ER against phosphatidylcholine (D’Souza et al., 2020). Unlike phosphatidylserine, phosphatidylcholine is not a limiting lipid species at the ER. Moreover, unlike other ORPs at the plasma membrane, the PH domain of ORP3 was reported to bind primarily PI(4,5)P2 (D’Souza et al., 2020) and could therefore function as a PI(4,5)P2 sensor. We hypothesized that ORP3 ability to bind plasma membrane PI(4,5)P2 and to transfer its PI4P precursor could allow to regulate both PI4P and PI(4,5)P2 at the plasma membrane.

ORP3 is involved in complex cellular processes dependent on PI(4,5)P2, such as cell adhesion, migration, and actin dynamics (Lehto et al., 2005; Lehto et al., 2008; D’Souza et al., 2020). Recent studies have linked ORP3 misregulation with aneuploidy and cancer development (Njeru et al., 2020; Wang et al., 2023; Xu et al., 2020). Thus, we investigated the potential role of ORP3 in PI(4,5)P2 regulation at the plasma membrane in mitosis.

Our results in HeLa cells demonstrate that ORP3 is strongly recruited by VAPA to the ER and to ER-plasma membrane contacts throughout mitosis and required for PI(4,5)P2 homeostasis. ORP3 is also required for PI4P extraction from the plasma membrane at the cytoplasmic bridge in abscission. ORP3 depletion or inactivation results in abnormal PI(4,5)P2, PI4P, cortical actin cytoskeleton distribution at the plasma membrane, leading to severe division defects throughout mitosis.

## RESULTS

### ORP3 inactivation impairs chromosome segregation during mitosis and cell division

In order to study ORP3 function in mitosis, we have established ORP3-KO HeLa cell lines, using a single nickase CRISPR-Cas9 strategy. Scrambled guide-RNAs were used to generate control CRISPR-Mock HeLa cell lines. Several independent clones were selected and tested by immunofluorescence and Western blotting (Fig. 1A, B). We confirmed that ORP3 was still present in CRISPR-Mock cells, where it is mostly cytosolic, similar to wild-type (WT) cells, but absent in the ORP3-KO cell lines. As recently reported (Wang et al., 2023), we observed that ORP3 depletion resulted in a significant increase of multinucleated cells, compared to WT or Mock cell lines in 5 different ORP3-KO clones tested (Fig. 1 C, D). Based on the consistent phenotype between various clones, only clone ORP3-KO1 was used in most of the following experiments. Using live cell imaging, we also observed that ORP3-KO cells exhibited various chromosome segregation defects (Fig. 1E) during. Only ∼ 40% of ORP3-KO cells in the population showed no defect, with chromosome segregation proceeding as in control cells (Fig. 1E WT, vs KO line 1a), while other cells presented various defects ranging from lagging chromosomes (Fig. 1E KO line b), chromosomes remaining trapped in the cytoplasmic bridge (Fig. 1E, line c), or chromosomes segregated to only one of the two daughter cells (Fig. 1E, line d). Additionally, a phenotype of abscission failure, where the two daughter cells re-merged at the end of cytokinesis, was observed in some ORP3 KO cells (Movie 1). Altogether, one or more division defect was detected in ∼ 60% of ORP3-KO cells, a value ∼ 6-fold higher than in WT or mock cells (Fig. 1F). Some of these defects resulted in cell aneuploidy, and eventually to the accumulation of multinucleated cells, as seen in Fig. 1C. These defects are likely to activate the spindle assembly checkpoint and to result in mitotic delays. In WT and Mock cells, the duration of mitosis, between nuclear envelope breakdown and the beginning of DNA decondensation after chromosome segregation, presented a unimodal distribution with times ranging from 45 to 75 minutes (Fig. 1G). ORP3 KO cells were distributed into 3 distinct populations. 48% had a profile similar to WT cells, 52% exhibited division times between 75 and 115 min (class II 23%), or times between 115 and 135 min (class III 29%), almost doubling the duration of mitosis. To determine if the division phenotypes observed in ORP3-KO cells were a primary consequences of ORP3 depletion, as opposed to indirect and/or cumulative defects in stable KO cell lines, we performed a transient ORP3 depletion by siRNA treatment of WT HeLa cells. Two distinct siRNAs resulted in a decrease in ORP3 protein signal in immunofluorescence and Western blots (Fig. 1H, I). Like ORP3-KO cells, ORP3 siRNA-treated cells exhibited various division defects, with a significant increase in frequency compared to scrambled control siRNA (Fig. 1J). Similar to ORP3-KO cells, ∼ 50% of cells showed at least one defect. In summary, our results show that ORP3 depletion results in chromosome segregation defects and delayed mitosis in more than 50% of cell divisions, and ∼15% multinucleated cells in the population at steady state. We hypothesize that these cells are eventually eliminated, resulting in a constant ploidy in the ORP3 KO cell population with no significant accumulation of defects in the established cell lines.

**Figure 1.**
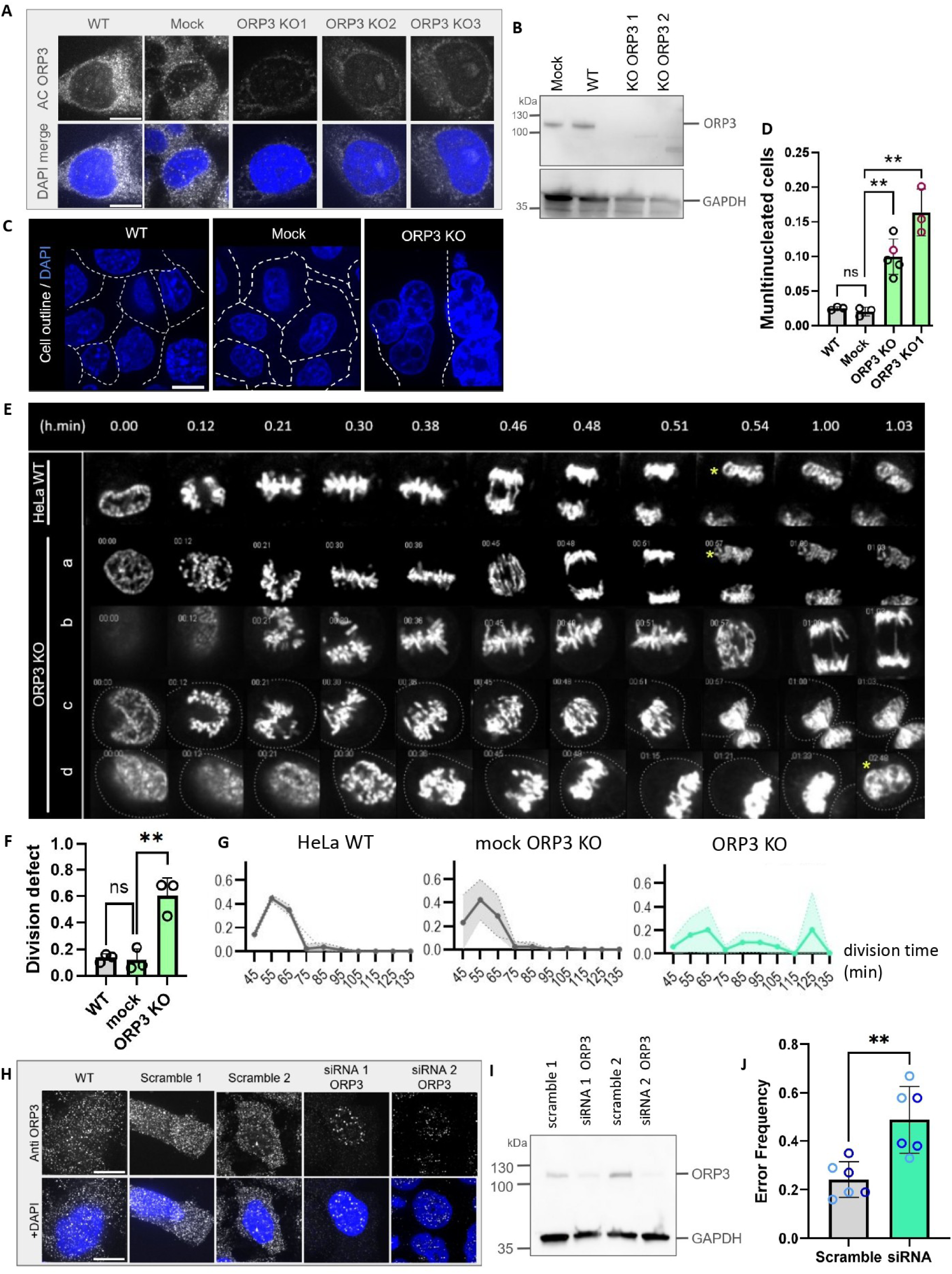
ORP3 is required for proper cell division. A Immunofluorescence staining with polyclonal anti-ORP3 polyclonal antibodies (white) in clonal cell lines from CRISPR Mock and ORP3-KO (3 distinct clones) HeLa cells. Nuclei were counter-stained with DAPI. Bar=10 µm. B Immunoblot staining for ORP3 from WT and CRISPR Mock or ORP3-KO cells (clones 1, 2). GAPDH was used as a loading control. C In fixed cells stained with DAPI, ORP3-KO cells (clone 1 shown) exhibited an accumulation of multinucleated cells, compared to WT or CRISPR Mock. Bar=20 µm. D Statistical analysis of the fraction of multinucleated cells in WT, Mock (3 distinct clones) and ORP3-KO cells (5 distinct clones). ORP3-KO Clone 1 (red circles), was used for the following experiments. E Live-cell imaging of mitosis from prophase to cytokinesis in WT HeLa cells (top) and ORP3-KO cells (lines a-d), showing typical ORP3-KO mitotic phenotypes: (a) no defect, (b) lagging chromosomes, (c) no DNA segregation between the two daughter cells and (d) segregation of all the DNA into only one of the daughter cells. DNA was stained with SiR-DNA. *: start of DNA decondensation. F Quantification of the total frequency of division defects in WT, Mock and ORP3 KO cell lines n=3 experiments. G Duration of cell division from nuclear envelope breakdown to the beginning of DNA decondensation in WT, Mock and ORP3-KO cells. The acquisition frequency was 3 minutes (10 minutes binning on the graph) n=3, 20-30 cells/experiment. H Immunofluorescence staining with anti ORP3 antibodies (white) and DAPI (blue) of WT, scramble siRNA- or ORP3 siRNA-treated HeLa cells. Bar=10 µm. I Immunoblot staining for ORP3 from WT, scramble siRNA or ORP3 siRNA-treated (2 distinct siRNAs) HeLa cells. GAPDH was used as a loading control. J Quantification of the division defects in scramble and ORP3 siRNA treated HeLa cells. Light blue: siRNA/Scr1, dark blue: siRNA/Scr2 (n=6).

### The Lipid transfer and membrane localization domains of ORP3 are all necessary for its function in mitosis

ORP3 functions at membrane contact sites between the ER - where it binds VAP proteins through a canonical FFAT motif - and the plasma membrane - where it binds PI(4,5)P2 through its “Plextrin Homology” (PH) domain (Fig. 2A). The ORD domain of ORP3 can transport PI4P from the plasma membrane to the ER and counter-transport phosphatidylcholine (D’Souza et al., 2020; Gulyás et al., 2020). ORP3 harbors an additional non-canonical FFAT-like motif, which requires phosphorylation to become active (see Fig. 4D) (Weber-Boyvat et al., 2015). This FFAT-like motif was hypothesized to be involved in the regulation of ORP3, but its contribution to ORP3 function remains unknown. To assess the importance of ORP3 domains to cell division, we performed rescue experiments in ORP3-KO cells. Stable HeLa ORP3-KO cells were stably transfected with either wild-type GFP-ORP3, or the different GFP-ORP3 mutants shown in Fig. 2B. In interphase, the six GFP-ORP3 mutants were mostly cytosolic, just like endogenous ORP3 (Fig. 2C). The expression of a protein of expected size at expression levels comparable to WT cells was confirmed by Western blotting using anti-ORP3 antibodies (Fig. 2D). We used the frequency of multinucleated cells, which was more than 15% in ORP3-KO cells, as a readout for the rescue of ORP3 function (Fig. 2E). As expected, Mock-KO HeLa had a low frequency of multinucleated cells, similar to WT cells. ORP3-KO cells stably expressing GFP-WT-ORP3 corrected the frequency of multinucleated cells, indicating that GFP-WT-ORP3 expression in ORP3-KO cells fully rescued this phenotype. This result also confirms that the mutinucleated phetonytpe is a direct consequence of ORP3 depletion. In contrast, stable cell lines expressing deletions of, or mutations in, any of PH, FFAT-like, FFAT motifs or ORD domains of ORP3 all exhibited elevated frequencies of multinucleated cells, with no or limited rescue of the ORP3-KO phenotype. These results strongly suggest that both the lipid transfer function of ORP3 and its localization at ER-plasma membrane contacts are necessary for proper cell division. In addition, our results also demonstrate that the often-overlooked FFAT-like motif is also essential to ORP3 function in mitosis. This is consistent with the idea that ORP3 can be “activated” through this domain, as previously proposed in response to PLC signaling (Gulyás et al., 2020). Deletion of the DLSK sequence of the ORD domain (Δ538-542), located in the lid of the hydrophobic pocket of ORP3 (Tong et al., 2021), further demonstrates that the transfer of lipid is also essential for proper division, beyond the mere presence of the ORD domain.

**Figure 2.**
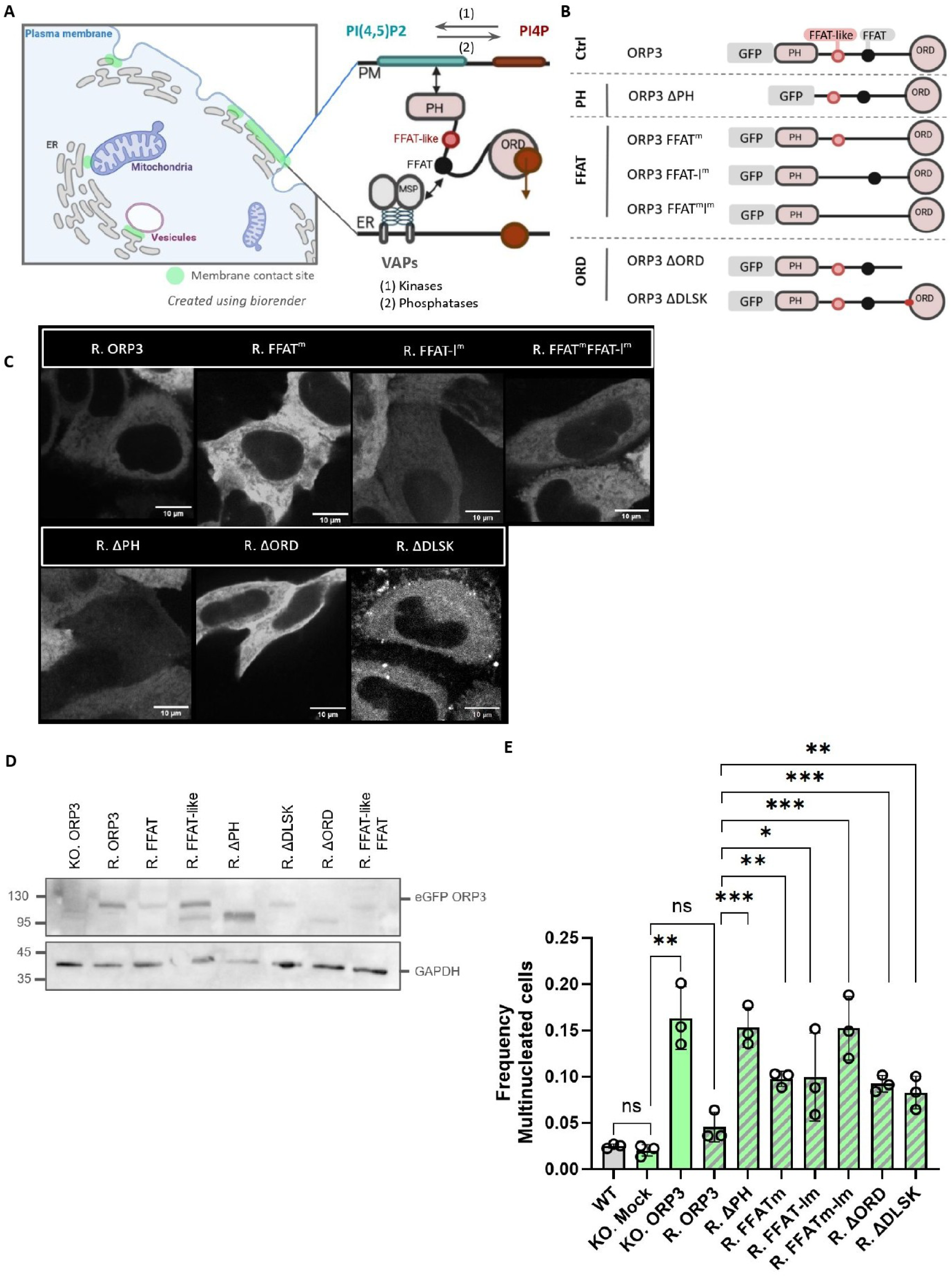
The major domains of ORP3 are necessary for its function in mitosis. A Scheme of ORP3 localization and function at membrane contact sites (light green) between the plasma membrane and the ER. B Scheme of the different GFP-ORP3 mutants used for rescue experiments in ORP3-KO HeLa cells. Mutations in the PH and ORD domains were deletions, whereas FFAT mutations were substitutions in the motif. C Localization of ORP3-KO HeLa cells expressing WT-GFP-ORP3 (R. ORP3) and mutants in live cells by spinning disk confocal microscopy. Bar=10µm. D Immunoblot staining with anti-ORP3 antibodies in GFP-ORP3 and rescue mutants stably expressed in ORP3-KO HeLa cells. Staining with anti-GAPDH antibodies was used as a loading control. Deletion of the PH (Δ1-150) or the ORD (Δ480-887) domain result in a smaller protein. E Frequency of multinucleated cells in WT cells, ORP3-KO, CRISPR Mock and ORP3 or mutated -ORP3 rescue cell lines, n=3 experiments for each condition.

### ORP3 is dramatically recruited to the ER by VAPA in mitosis

The ORP3 localization domains are required for cell division, yet, as shown in Fig. 2C and Fig. 3A (top inset), ORP3 is mostly cytosolic in interphase and only weakly ER- or plasma membrane-associated. Therefore, we investigated the localization of ORP3 during mitosis in ORP3-KO HeLa cells stably expressing GFP-ORP3 by live-cell imaging in spinning disk confocal microscopy (Fig. 3A). From prometaphase, when the ER starts tubulating, until telophase (Puhka et al., 2007), ORP3 exhibited a dramatic change in subcellular localization, from primarily diffuse to a network pattern, resembling the ER. To confirm that ORP3 is associated with the ER in mitosis, we co-expressed WT GFP-ORP3 and mCherry-VAPA for live-cell imaging in ORP3-KO HeLa cells (Fig. 3C, top - metaphase shown). These proteins extensively co-localized, indicating that ORP3 strongly localizes to the ER during mitosis. ORP3 was also heavily present at the cell cortex and poles (Fig. 3A), suggesting that it could function at ER-plasma membrane contacts all along mitosis. In late cytokinesis, prior to abscission, ORP3 staining was restored to a diffuse localization in the cytosol, in addition to a strong enrichment at the midbody (Fig. 3A, bottom inset). The presence of ER-plasma membrane contacts in mitosis has been debated (Yu and Machaca 2025). We have confirmed their existence using the ER-plasma membrane contacts probe GFP-MAPPER (Chang et al., 2013), which showed a dense punctate staining at the plasma membrane throughout mitosis (Fig. 3B). ER-plasma membrane contacts were also observed during mitosis by electron microscopy, even in ORP3-KO cells (Fig. 3E). In addition, WT-ORP3 (Fig. 3C, left panels) exhibited a punctate staining at the cell cortex, reminiscent of the GFP-MAPPER pattern. These results indicate that ORP3 is recruited to the ER throughout mitosis and is probably also located at ER-plasma membrane contacts.

**Figure 3.**
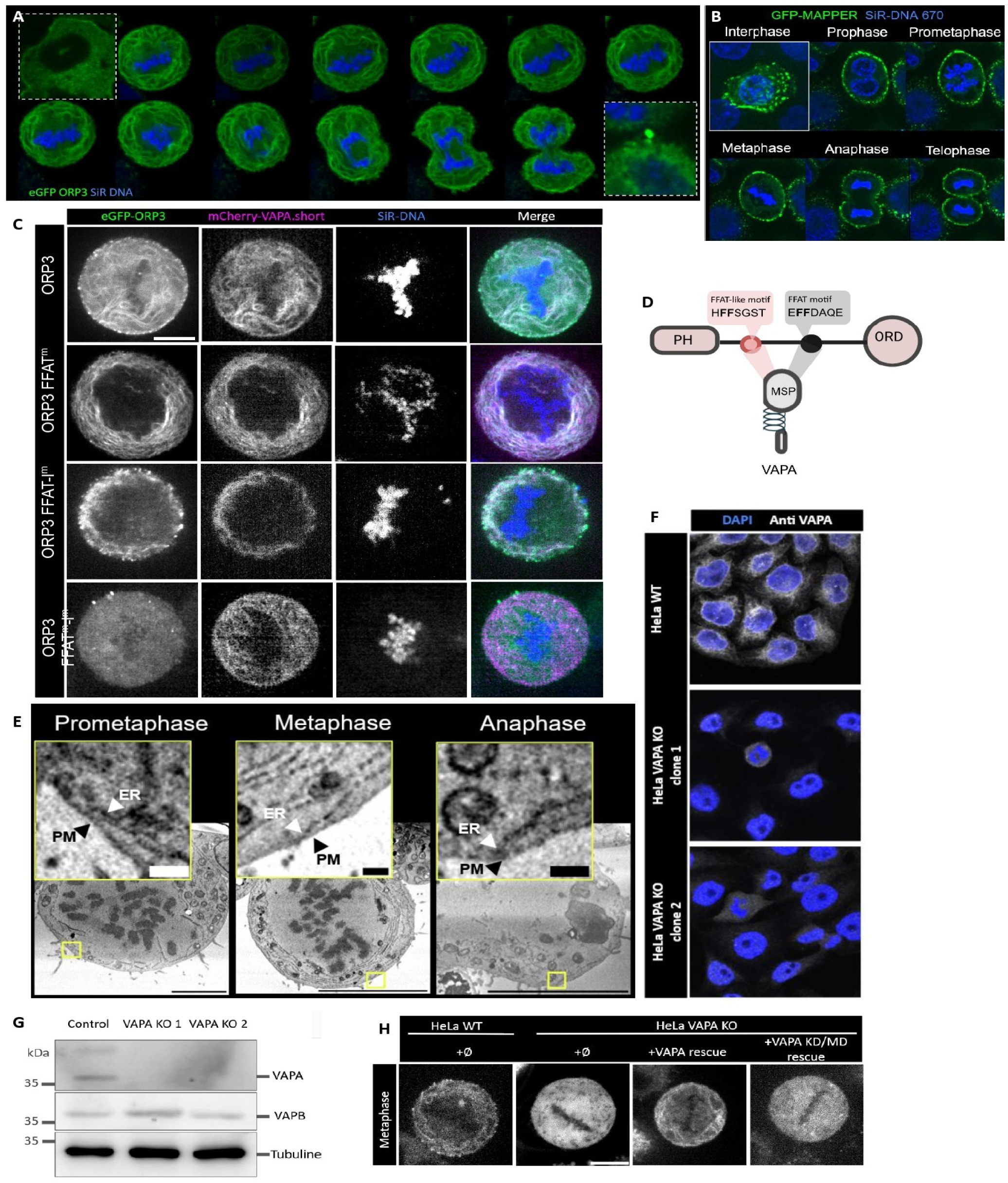
ORP3 is strongly recruited to the ER by VAPA through both ORP3 FFAT motifs. A Live-cell imaging of stable GFP-ORP3 HeLa cells stained with SiR-DNA (blue) during mitosis. Top-left inset: in interphase, ORP3 is mostly cytosolic. ORP3 is recruited to the ER from prophase to telophase, with a final enrichment at the midbody later in abscission (bottom-right inset). B localization of the ER-plasma membrane contact probe GFP-MAPPER in interphase (inset) and throughout mitosis. Blue: Sir-DNA staining. C FFAT-dependent ER localization of ORP3. In live cell imaging, ORP3-KO HeLa cells stably expressing WT-GFP-ORP3 co-localizes with mCherry-VAPA at the ER. This localization persists if either FFAT (ORP3-FFAT^m^) or FFAT-like (ORP3-FFAT-like^m^) are inactivated, but not when both motifs are defective. D Detailed scheme of ORP3 FFAT and FFAT-like motif sequence. E SBF-SEM electron microscopy of ORP3-KO HeLa cells confirms the existence of ER-plasma membrane contacts from metaphase to anaphase even in the absence of ORP3. F Anti-VAPA antibody staining (white) in WT HeLa cells and stable VAPA-KO cell lines. DNA is stained with DAPI (blue). G Immunoblot staining of VAPA and VAPB in control and VAPA-KO HeLa cell lines. Only the VAPA signal is abolished in VAPA-KO cells. Tubulin was used as loading control. H VAPA-dependent ER localization of ORP3. Live-cell imaging in metaphase of stable GFP-ORP3 in WT-HeLa cells (left) and VAPA-KO HeLa cells (second from left), VAPA-KO cells rescued with WT-VAPA (third from left) or VAPA mutated in its FFAT-binding MSP domain (VAPA KD/MD, right). Bar=20µm.

We next investigated the molecular basis for ORP3 localization to the ER during mitosis. First, we examined whether the FFAT motifs of ORP3 (FFAT and FFAT-like) were involved in ORP3 localization during mitosis (Fig. 3D pink and grey respectively). As shown in Figure 3C, mutation in either one of the two FFAT motifs (FFAT^m^ and FFAT-like^m^) was not sufficient to delocalize ORP3 from the ER in mitosis (Fig. 3C, lines 2 and 3). However, in double FFAT^m^-FFAT-like^m^ mutants, ORP3 became totally cytosolic (Fig. 3C, bottom line). These results demonstrate that both the canonical FFAT motif (as expected) and non-canonical FFAT-like motif contribute to ER localization of ORP3 during mitosis. We then examined whether the ER localization of ORP3 was dependent on its ER anchoring partners, the VAP proteins. In HeLa cells, VAPA is the most abundant VAP (Di Mattia et al., 2018). A VAPA-KO HeLa cell line was created using a double nickase CRISPR-Cas9 strategy. Several clones were produced (two shown). The VAPA signal was abolished both in immunofluorescence (Fig. 3F) and in Western blotting (Fig. 3G). Importantly, VAPA deletion did not change the expression of the less abundant VAPB protein (Fig. 3G). In VAPA-KO HeLa cells, WT-ORP3 was no longer recruited to the ER in mitosis (Fig. 3H). Transfection of VAPA-KO cells with a CRISPR-resistant VAPA rescued the ORP3 localization, but not transfection with a “dead” VAPA, mutated in its MSP domain of interaction with FFAT motifs (Fig. 3H, VAPA KD/MD). We conclude that during mitosis, ORP3 is strongly recruited by the MSP domain of VAPA to the ER through both its canonical (FFAT) and non-canonical (FFAT-like) motifs.

### The recruitment of ORP3 by VAPA to the ER is enhanced by phosphorylation of the ORP3 FFAT-like motif

The interaction between VAPA and ORP3 in interphase and mitosis was further investigated by the study of ORP3 co-immunoprecipitation with VAPA, using a double thymidine nocodazole block to synchronize cells in G2/M (Fig. 4A). The abundance of VAPA in cells was not changed by the block, nor was the total abundance of ORP3 (Fig. 4A, input). However, there was a shift in the migration of ORP3 bands in synchronized cells, suggesting a post-translational modification of ORP3 at the onset of mitosis. In both non-synchronized and synchronized cells, ORP3 was present in VAPA immunoprecipitates (Fig 4A, VAPA-IP). However, in addition to the migration shift, the amount of immunoprecipitated ORP3 was much higher in synchronized cells, suggesting a strong increase in the interaction between VAPA and ORP3 in prometaphase. This result is consistent with the ORP3 recruitment to the ER observed in live-cell imaging (Fig. 3C). We hypothesized that the FFAT-like motif of ORP3 was activated during mitosis, *i.e.* phosphorylated to create an acidic-like domain (Weber-Boyvat et al., 2015) and that the post-translational modification of ORP3 in mitosis was due to phosphorylation. Indeed, a very significant part of the proteome (∼30%) is phosphorylated in mitosis. After treatment of cell extracts from synchronized cells with the CIP phosphatase (Fig. 4B), the migration shift of ORP3 was abolished confirming that ORP3 is phosphorylated in mitosis. Considering the large change in migration speed, this result suggests that a number of residues of ORP3 are phosphorylated during mitosis. We used mass-spectrometry to establish a map of phosphorylated residues in ORP3 in synchronized cells. As expected, the FFAT-like motif itself, as well as several amino acids downstream, were selectively phosphorylated in mitosis (Fig 4C, yellow residues). To investigate further the consequences of ORP3 FFAT-like motif activation by phosphorylation, we constructed a phosphomimetic mutant of the ORP3 FFAT-like motif, by substitution of residues corresponding to acidic residues in the canonical FFAT motif, not including serines downstream (Fig. 4D, green residues). We then examined the localization of this “phospho-FFAT-like” mutant in transiently-transfected HeLa cells, compared to WT-ORP3 (Fig. 4E, F). As we expected, this mutant was more strongly localized to the ER than WT-ORP3 even in interphase, particularly around the nuclear membrane and in the vicinity of the Golgi apparatus where ER is abundant. As we often observed in transient ORP3 transfections, the localization of phosphomimetic mutant ORP3 to the plasma membrane was also enhanced, thus suggesting that a very significant part of ORP3 is located at ER-plasma membrane contacts. Transfected cells were not observed to enter mitosis and died rapidly, suggesting that ORP3 recruitment to contact sites needs to be finely regulated in the cell cycle. To confirm that the ER localization of the phospho-FFAT-like ORP3 mutant was mediated by VAPA, we transfected VAPA-KO Hela cells with this mutant (Fig. 4G, H). As expected, the ER localization of the phospho-FFAT-like mutant was strongly decreased - but not fully abolished. It is possible that a fraction of the protein, which has a higher affinity for the VAP MSP domain, is recruited by VAPB, the less abundant VAP in HeLa cells. From Figures 3 and 4, we conclude that the ORP3 FFAT-like motif is phosphorylated in mitosis and that this phosphorylation enhances the recruitment of ORP3 by VAPA to the ER between prophase and cytokinesis.

**Figure 4.**
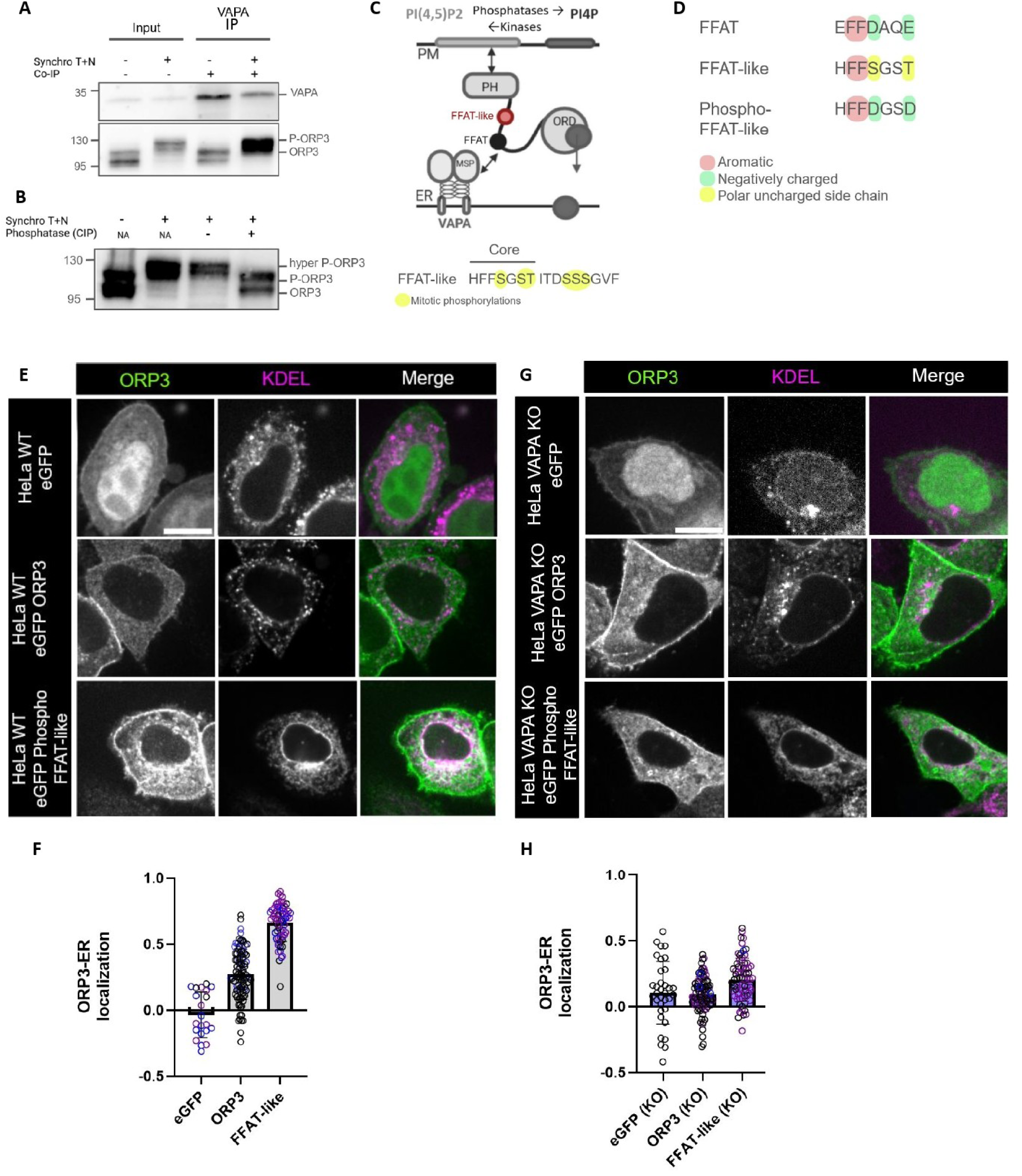
Phosphorylation of the FFAT-like motif of ORP3 enhances its ER localization. A VAPA-immunoprecipitation of non-synchronized HeLa cells and cells synchronized by a double thymidine + nocodazole block (Synchro T+N). VAPA is detected in the input (left), and after VAP immunoprecipitation (VAPA IP) in both conditions. ORP3 is present in the input in comparable amounts but display a migration delay in synchronized cells. In synchronized cells, ORP3 co-immunoprecipitation with VAPA is strongly enhanced. B In HeLa cells extracts from synchronized (Synchro T+N) or non-synchronized, treatment with the CIP phosphatase reverses the migration delay of ORP3 in synchronized cells. The left two lanes (NA) were not subject to the enzymatic incubation. C Scheme of ORP3 and the sequence of its FFAT-like motif (Core) and amino acids downstream. Residues highligthed in yellow were found phoshorylated in synchronized cells, but not non-synchronized cell by mass spectrometry, including the three S and T residues of the FFAT-like motif and 3 serines downstream. D Design of a FFAT-like phosphomimetic mutant, where S and T residues in the FFAT-like motif, which correspond to acidic residues in canonical FFAT, were replaced by aspartic acid (D). E WT-HeLa cells double-transfected with mCherry-KDEL and GFP, WT-GFP-ORP3 or the FFAT-like phosphomimetic mutant of ORP3 (GFP-phospho-FFAT-like). Bar=20µm. F Quantification of the ER co-localization of GFP alone, GFP-WT-ORP3, and GFP-phosphomimetic mutant (FFAT-like). G VAPA-KO HeLa cells double-transfected with mCherry-KDEL and GFP, WT-GFP-ORP3 or the FFAT-like phosphomimetic mutant of ORP3 (GFP-phospho-FFAT-like). Bar=20µm - H Quantification of the ER co-localization of GFP alone, GFP-WT-ORP3, and GFP-phosphomimetic mutant (P-FFAT-like) in VAPA-KO HeLa cells.

### ORP3 regulates PI4P and PI(4,5)P2 at the plasma membrane in mitosis

To understand the molecular mechanism behind the multinucleated phenotype of ORP3-KO cells, we studied the contribution of ORP3 to the distribution of PI4P and PI(4,5)P2. Compared to interphase, it is well established that PI(4,5)P2 strongly accumulates at the plasma membrane in mitosis (Roubinet et al., 2011). In addition, PI(4,5)P2 distribution at the plasma membrane varies along mitosis, with an accumulation at the mitotic furrow in between anaphase and cytokinesis, in order to facilitate the recruitment of actin for ingression of the furrow (Logan and Mandato 2006; Cauvin and Echard 2015). In cells stably expressing a fluorescent PI(4,5)P2 probe (mCherry-PH-PLCẟ), the fluorescent signal was much stronger in mitosis from metaphase to late anaphase (Fig. 5A) than in interphase. The PI(4,5)P2 probe signal was homogeneous along the cell cortex in metaphase, but there was a progressive increase of the relative PI(4,5)P2 signal at the ingression furrow from metaphase to late cytokinesis (Fig. 5A,B). In WT cells, a 60% increase in the PI(4,5)P2 signal was measured at the furrow (corresponding to x-axis position 25 and 75 along the cell contour on Fig. 5B, grey line) compared to the poles (Fig. 5B, grey baseline, Fig. 5C). Several phenotypes were observed in ORP3-KO cells. In some cells, the PI(4,5)P2 distribution was asymmetric at the furrow (Fig. 5A a, b). In others, the PI(4,5)P2 increase extended well beyond the furrow (Fig. 5Ab). In many cases, the size of two future daughter cells was different (Fig. 5 Ab). In ORP3-KO cells, the PI(4,5)P2 signal was very strong at the plasma membrane, but the ratio of PI(4,5)P2 at the furrow to the poles culminated at only 30% in cytokinesis (Figs. 5B, 5C green). The PI(4,5)P2 intensity peaks at the furrow in cytokinesis were also broader in ORP3-KO cells, suggesting impaired PI(4,5)P2 accumulation at the ingression furrow. Due to inter-cellular differences in expression levels of the PH-PLCẟ probe and signal intensity in non-clonal cell lines, the absolute PI(4,5)P2 concentration could not reliably be measured and compared between the two conditions. Nevertheless, our results demonstrate that ORP3 deletion results in impaired PI(4,5)P2 distribution at the plasma membrane during mitosis, and altered symmetry of the equatorial plane at the onset of cytokinesis. In HeLa cells expressing endogenous ORP3, PI4P was not particularly enriched at the plasma membrane during early mitosis (not shown). This is consistent with the strong activation of PI4P-5K kinases, which convert PI4P into PI(4,5)P2 (Roubinet et al., 2011). In abscission however, we observed a strong PI4P signal at the midbody in WT cells (Fig. 5D, top), where ORP3 is also particularly abundant (see Fig. 3A, bottom inset). Cells overexpressing GFP-ORP3-ΔORD (unable to extract PI4P) presented a dramatic increase in PI4P all along the cytoplasmic bridge (Fig. 5D, bottom). This phenotype demonstrates the requirement for ORP3 PI4P transfer function at the bridge. Abscission requires PI(4,5)P2 depletion at the bridge by the OCRL phosphatase (Dambournet et al., 2011). Since OCRL catalyzes the hydrolysis of PI(4,5)P2 into PI4P, our results show that ORP3 is required downstream of OCRL to drain PI4P from the plasma membrane and facilitate PI(4,5)P2 hydrolysis.

**Figure 5.**
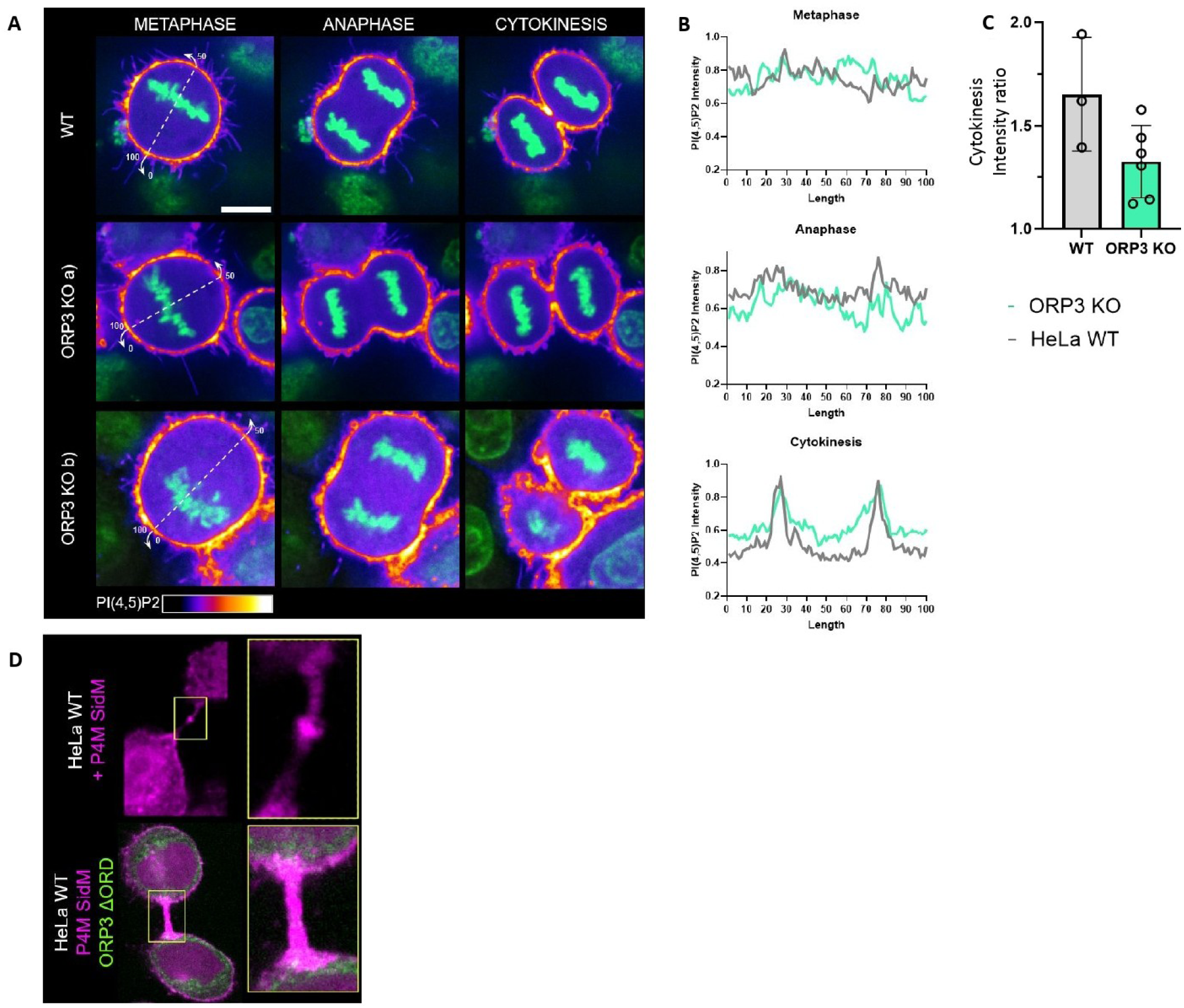
- ORP3 regulates PI4P and PI(4,5)P2 distribution at the plasma membrane during mitosis. A Live cell imaging spinning disk confocal microscopy of PI(4,5)P2 and DNA (SiR DNA 670, blue) at 3 different mitotic stages in WT and ORP3-KO cells stably expressing mCherry-PH-PLCẟ. Bar=20µm. PI(4,5)P2 intensity is represented by a heat-map LUT. B Quantification of the relative PI(4,5)P2 intensity along the plasma membrane in each of the 3 phases (arbitrary units). Membrane length was normalized with position 0/100 corresponding to one pole and 25 (and 75) to the equatorial plane. Grey line: WT, Green line: ORP3-KO cells. C Quantification of PI(4,5)P2 intensity at the furrow in cytokinesis compared to the poles. n=2 experiments, each dot corresponds to one cell. D Live cell imaging spinning disk confocal microscopy of PI4P (mCherry-P4MSiDM) during cytokinesis in WT HeLa cells (top), or cells expressing GFP-ORP3-ΔORD (green, bottom).

### Cortical actin and the mitotic spindle are dramatically altered in ORP3-KO cells during mitosis

In view of the alterations in PI(4,5)P2 distribution at the plasma membrane in ORP3-KO cells, we next investigated the consequences of ORP3 deletion on cortical F-actin, which is heavily dependent on PI(4,5)P2. Cortical actin staining intensity was measured in live-cell imaging using spinning-disk microscopy and the SiR-actin probe at different stages of mitosis (Fig. 6A). Unlike the PI(4,5)P2 probe, the SiR-actin probe gave a very reproducible intensity readout in each of the two conditions. The actin staining was much more intense in ORP3-KO cells compared to WT cells. The average cortical actin intensity signal was therefore measured and compared between WT and ORP3-KO cells at different stages of mitosis (Fig. 6B). A striking and very significative ∼4 fold increase in cortical actin was observed in ORP3-KO cells compared to controls from prophase to telophase. This result demonstrates a dramatic thickening of cortical F-actin in mitosis in the absence of ORP3, consistent with an overall upregulation of PI(4,5)P2. The actin intensity profile along the cell cortex was measured in cytokinesis (Fig. 6C). In addition to a markedly elevated average actin intensity, the accumulation of actin at the ingression furrow (relative positions 25 and 75, poles at 0 and 100) was even more dramatically upregulated in ORP3-KO HeLa cells, compared to WT cells. From our results on PI(4,5)P2 and actin distribution, we propose that ORP3 deletion results in abnormal PI(4,5)P2 accumulation and distribution at the plasma membrane, a massive and aberrant increase in cortical actin recruitment in mitosis. PI(4,5)P2 and cortical actin are also important determinants of the assembly and positioning of the mitotic spindle. Using SIR-tubulin staining of microtubules in live cell imaging, we investigated the spindle morphology in metaphase (Fig. 6D). A statistical analysis of the spindle length and width (Fig. 6D horizontal and vertical dotted lines) revealed that the spindle width was reduced in ORP3-KO cells compared to WT cells, whereas no difference was detected in spindle length (Figs. 6E, 6F). Thus ORP3 deletion, in addition to abnormal cortical actin distribution, results in abnormal spindle morphology.

**Figure 6.**
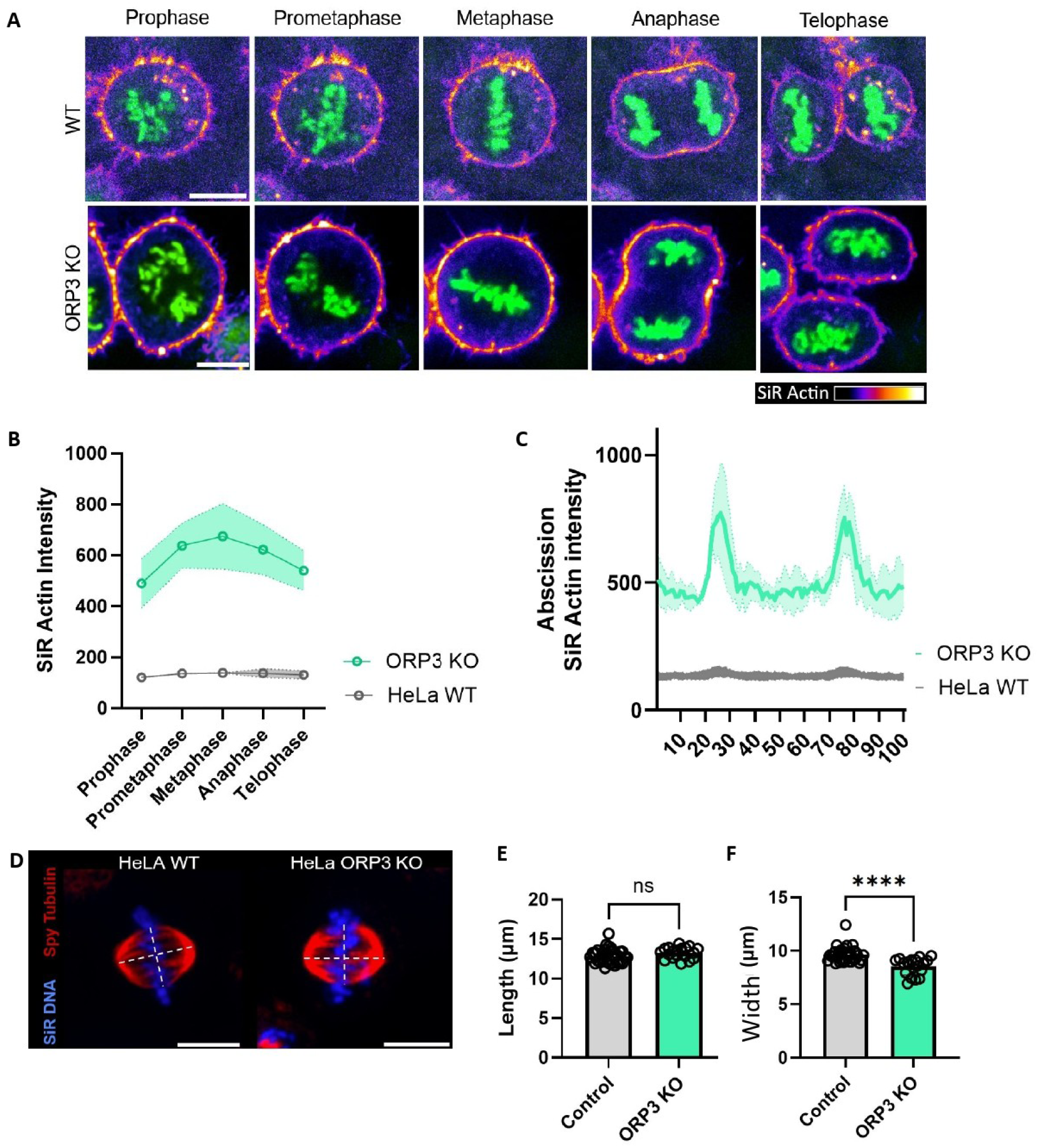
ORP3 is required for actin and tubulin distribution in mitosis. A Live cell imaging spinning disk confocal microscopy of actin (Sir Actin) and DNA (Spy-DNA, green). Bar = 20µm. B - Quantification of the average actin signal at the cortex across mitotic phases. Shades: S.E.M. C Quantification of the actin signal intensity along the cell cortex in cytokinesis. Membrane length was normalized with position 0/100 corresponding to each pole and 25 (and 75) to the equatorial plane. Shades: S.E.M. D Fluorescence staining of the mitotic spindle in metaphase in WT and ORP3 KO cells (red, Tubulin, Blue DNA). E Length of the mitotic furrow in control and ORP3 KO cell lines (n=3). F Width of the mitotic furrow in control and ORP3 KO cell lines (n=3).

## DISCUSSION

The lipid composition of cell membranes contributes to defining the membrane identity and functions. In mitosis, PI4P-5K kinases are strongly activated and the increase in PI(4,5)P2 that results is essential for thickening of cortical F-actin (Kunda et al., 2008) and for the anchoring on the mitotic spindle (Machicoane et al., 2014; Kotak et al., 2014). In addition to vesicular trafficking and to the activity of lipid-modifying enzmes, direct transfer of lipids, including phosphoinositides, at membrane contact sites have emerged as a key mechanism to regulate the lipid composition of membrane organelles.

In the present work, we propose that ORP3, a lipid transfer protein of the ORP family, is essential to the regulation of PI(4,5)P2 and PI4P at the plasma membrane during mitosis.

A common feature of many ORP lipid transfer proteins (OSBP, ORP5, ORP8) is to use PI4P transfer as a driving force for the counter-transport of a different lipid species against its gradient (Mesmin et al., 2013; Moser von Filseck et al., 2014; Antonny et al., 2018). Their PH domain also binds PI4P, which contributes to their downregulation when PI4P concentrations decrease. In this respect, ORP3 stands out as an exception. ORP3 also transports PI4P but it binds primarily PI(4,5)P2 at the plasma membrane (D’Souza et al., 2020; Gulyás et al., 2020). The PH domain of ORP3 can thus act as a PI(4,5)P2 sensor (Fig. 7).

**Figure 7.**
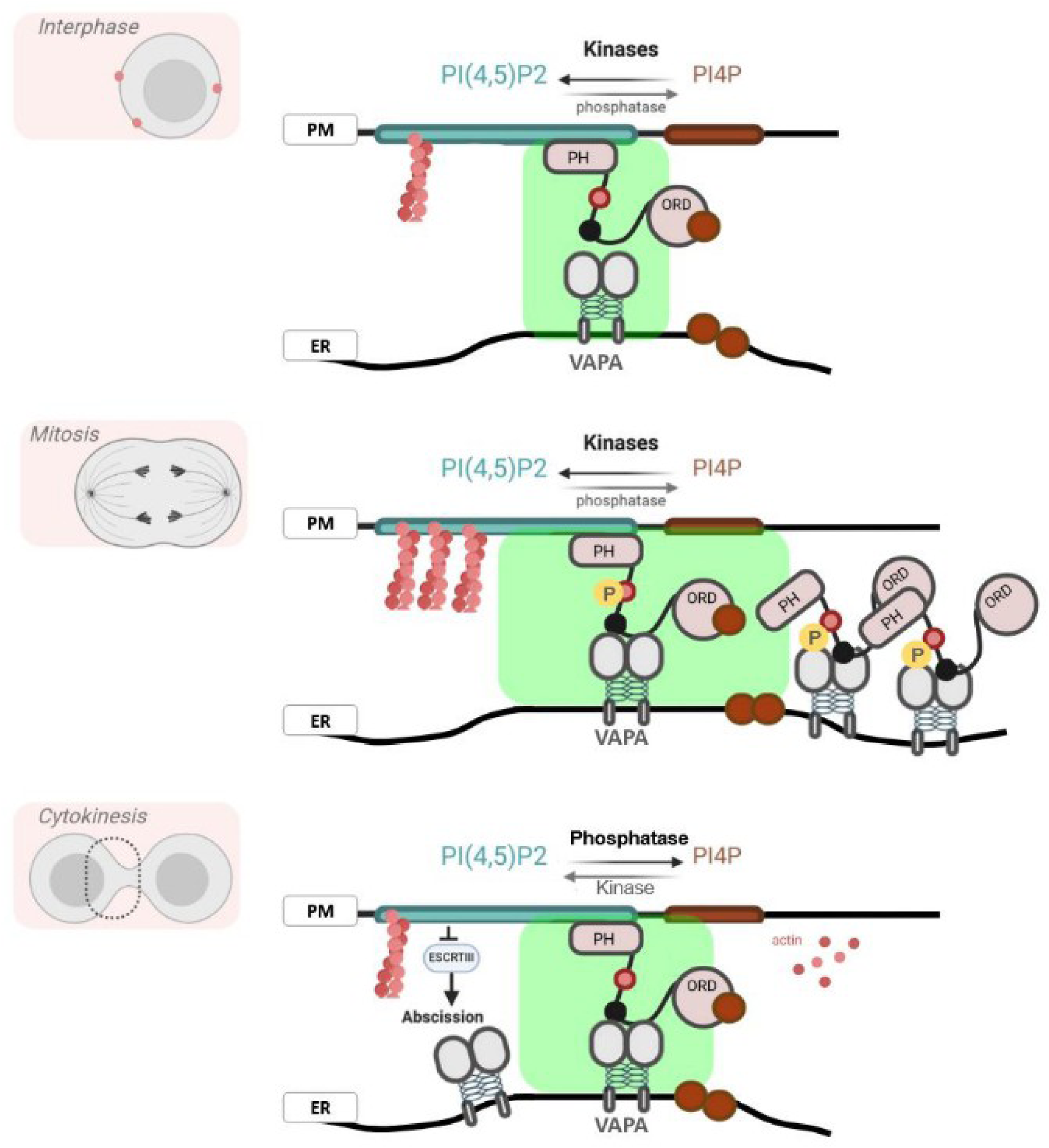
Proposed function and regulation of ORP3 regulation in the cell cycle. In interphase (top panel), ORP3 is associated to PI(4,5)P2-rich plasma membrane domains and only weakly to the ER. ORP3 interaction domains as well as the inter-conversion between PI4P and PI(4,5)P2 by phosphatases and kinases can be regulated by intracellular signaling. During mitosis (middle panel), the ORP3 FFAT-like motif is phosphorylated and ER-association of ORP3 is strongly enhanced. Since plasma membrane PI(4,5)P2 levels are also increased at the plasma membrane, association of ORP3 at ER-plasma membrane contacts is potentiated. Following this ORP3 “priming” at contact sites, the regulation of PI4P and PI(4,5)P2 becomes highly dependent on local phosphatases and kinases. At the end of cytokinesis, ORP3 is associated to the cytoplasmic bridge, where PI(4,5)P2 levels are elevated. Phosphatases such as OCRL are activated in order to hydrolyze PI(4,5)P2 into PI4P and ORP3 function allows to drain and deplete PI4P from the plasma membrane at the cytoplasmic bridge and contribute to PI(4,5)P2 reduction.

In interphase, ORP3 is weakly and transiently associated with the ER, through its canonical FFAT motif, and with the plasma membrane through its PI(4,5)P2-binding PH domain (Fig. 7, top). ORP3 function is known to be activated by post-translational modifications in response to R-Ras and/or PLC signaling, which also affect PI(4,5)P2 at the plasma membrane (Weber-Boyvat et al., 2015). Here, we demonstrate that in mitosis (Fig. 7, middle panel), the non-canonical FFAT-like motif of ORP3 is phosphorylated and activated, which results in enhanced ORP3 association with VAPA at the ER. In addition, plasma membrane PI(4,5)P2 levels are elevated, also increasing plasma membrane association of ORP3. We propose that this results in “priming” of ORP3 at ER-plasma membrane contacts, where it is readily available for PIP4 transport. In concert with kinases and phosphatases, ORP3 can then contribute to the regulation of PI4P and PI(4,5)P2 at the plasma membrane. Finally, unlike other lipid transfer proteins, ORP3 was shown to counter-transport an abundant lipid: phosphatidylcholine (D’Souza et al., 2020) suggesting that PI4P can be the rate-limiting substrate of ORP3.

Altogether, we propose that ORP3 function and regulation must be investigated under a new perspective. When located at contact sites where PI(4,5)P2 is abundant ORP3 can effectively extract PI4P, instead of using it as a driving force, until PI4P is depleted or PI(4,5)P2 concentrations become low enough for ORP3 to detach.

Accordingly, our results demonstrate that ORP3 function is necessary throughout mitosis to maintain the PI(4,5)P2 distribution at the dividing cell poles and ingression furrow. In the absence of ORP3, PI(4,5)P2 distribution is impaired and a dramatic increase in cortical actin is observed all along the plasma membrane and even more so at the ingression furrow. Whether the spatial dynamics of ER-plasma membrane contact sites in mitosis is also regulated remains to be investigated.

Abnormal stiffening of the F-actin cortex was previously reported to impair cell elongation, contractility, and to disturb the assembly and positioning of the mitotic spindle (Roubinet et al., 2011). Interestingly, we found that the shape of the spindle was altered in ORP3-KO cells. In metaphase, the length of the spindle was normal, but its width was significantly reduced in ORP3-KO cells. It has been demonstrated that spindle length and width are coupled and that their scaling contributes to silencing of the spindle assembly checkpoint (Hara and Kimura, 2013; Chen and Liu, 2016), which underscores the importance of the spindle geometry in cell division. Consistent with the delayed mitosis phenotype, the abnormal spindle organization in ORP3-KO cells is likely to delay the spindle assembly checkpoint. Moreover, it was proposed that spindle width is a better readout of spindle volume than spindle length (Kletter *et al*., 2022). This suggests that the total volume of the spindle is decreased in ORP3-KO cells, which could lead to the chromosome segregation defects observed in ORP3-KO cells.

Finally, ORP3 function is absolutely essential to clear PI4P from the cytoplasmic bridge prior to abscission (Fig. 7, bottom). It is well established that the OCRL phosphatase is required to convert PI(4,5)P2 into PI4P and to lower PI(4,5)P2 levels at the bridge (Carim et al., 2019). This step is crucial for releasing the actin cytoskeleton and to permit the recruitment of ESCRT complex for abscission. We found that in the absence of a functional ORP3 protein, PI4P dramatically accumulates at the cytoplasmic bridge and it probably restricts PI(4,5)P2 depletion. To our knowledge, ER-plasma membrane contact sites, where ORP3 could function, were never described at the cytoplasmic bridge. However, the accumulation of ORP3 at the midbody in control cells strongly supports an ORP3 function at the abscission site. When non functional ORP3 is overexpressed, we see elevated PI4P levels at the bridge, which likely delays or impairs abscission. Indeed, abscission failure and daughter cells fusion after mitosis are on of the observed in ORP3-KO cells.

In conclusion, beyond the previously described role of ORP3 in interphase, we show that ORP3 transport of PI4P is essential for the exquisite regulation of PI(4,5)P2, PI4P, and the cytoskeleton at different plasma membrane domains throughout mitosis, and thus essential for cell division.

## ACKNOWLEDGMENTS

We warmly thank Arnaud Echard, Katja Wassmann, Fabien Alpy, Phong Tran and the past and current members of the Membrane Dynamics and Intracellular Trafficking team at Institut Jacques Monod for insightful discussions. We acknowledge the ImagoSeine core facility of the Institut Jacques Monod, member of the France BioImaging infrastructure (https://ror.org/01y7vt929) supported by the French National Research Agency (ANR-24-INBS-0005 FBI BIOGEN) and GIS-IbiSA and the support of the Région Île-de-France (Sesame) for flow cytometry, light microscopy, and electron microscopy. This work was supported by grants from the Ecole doctorale Bio Sorbonne Paris Cité and the Labex “Who Am I” (A.V), the Agence Nationale de la Recherche (ANR 225916 EndoMitR and ANR 237213 SynaptoLigation), the Ligue Contre le Cancer (RS21/75-85).

## MATERIALS AND METHODS

### Cell culture

Hela cell lines were cultured in Dulbecco’s modified Eagle’s medium (DMEM) 1g/L D-glucose medium (Gibco), supplemented with 10% fetal bovine serum (FBS, Gibco) and 100 units/ml penicillin (Gibco), in 5% CO_2_ at 37°C. HeLa AP cells were edited with a double nickase strategy to generate the CRISPR control and VAPA-KO cell lines (Siegfried et al., 2024). HeLa AP cells were edited with a single nickase strategy to generate the CRISPR control and ORP3-KO cell lines (cf “Cell line editing using CRISPR/Cas9” section). All stable cells lines (ORP3 KO-rescues) were grown in DMEM 1g/L D-glucose medium, supplemented with 10% FBS and 250 units/ml Geneticin G418 (Gibco).

### Cell transfection

Cells were transfected with Lipofectamine™ (Thermofisher). DNA Plasmid transfection was performed using Lipofectamine™ 2000 (Thermofisher), according to manufacturer’s recommandations. HeLa cells were transfected at a 60-70% confluence. For transfection, 1 µg plasmidic DNA was resuspended in 100µl OPTI MEM (1X) + GlutaMAX™-I media (Gibco). 3µl lipofectamine 2000 was resuspended in 100µl OPTI MEM. Resuspended DNA and lipofecamine 2000 were mixed for 10 min, then homogenized with the cell medium for transfection. 4h after transfection, the medium was replaced with DMEM supplemented with 10% FBS. Experiments on transfected cells were performed the next day. siRNA plasmid transfection was performed with Lipofectamine™ 3000 (Thermofisher) according to manufacturer’s instructions. HeLa cells were transfected at a 50-60% confluence. 2µg siRNA were used for 5µl of lipofectamine 3000. 2µg of siRNA was resuspended in 200µl OPTI MEM (1X) + GlutaMAX™-I media. 5µl lipofectamine 3000 was resuspended in 200µl OPTI MEM. Resuspended DNA and lipofectamine 2000 were mixed for 10 min, then homogenized with the cell medium for transfection. Cells were incubated overnight in the transfection medium. The medium was then replaced with DMEM containing FBS. Experiments were performed three days after transfection.

### Live-cell imaging probes

The Fluorescent probes used to stain cell compartments for live-cell imaging were diluted as much as possible (see below) to limit side-effects. SiR-DNA (λ674) (Spirochrome) and Spy-DNA (λ505) (Spirochrome) were prepared with 0,4 nM verapamil enhancer according to manufacturer’s instructions. They were diluted 1:2500 in FluoroBrite™ DMEM supplemented with 10% fetal bovine serum and 100 units/ml penicillin OPTI MEM. SiR-Actin (λ674) and Spy-Tubulin (λ505) were used at a 1:5000 dilution.

### Antibodies

Primary antibodies used for immunofluorescence and their dilutions: anti-ORP3 rabbit IgG polyclonal (1:300, Abcam, 224212); anti-PI4P mouse IgM monoclonal (1:200, Echelon Biosciences, Z-P004); anti-PI(4,5)P2 mouse IgM monoclonal (1:200, Echelon Biosciences, Z-P045). Secondary antibodies used for immunofluorescence: Alexa Fluor™ 488 Goat anti-Mouse IgG polyclonal (1:10000, Invitrogen, A11001); Alexa Fluor™ 488 Goat anti-Rabbit IgG polyclonal (1:10.000, Invitrogen, A11034); Alexa Fluor™ 555 Goat anti-Mouse IgM polyclonal (1:10.000, Invitrogen, A21426). Primary antibodies used for the western-blotting: anti-ORP3 mouse IgG monoclonal (1:1000, Santa Cruz Biotechnology, sc-398326); anti-VAPA rabbit polyclonal (1:1000, Atlas Antibodies, HPA009174); anti-GAPDH mouse IgG monoclonal (1:2000, GeneTex, GTX627408); anti α-tubulin mouse IgG monoclonal (1:1000, GeneTex, GTX628802). Secondary antibodies used for western-blotting: Peroxidase anti-mouse IgG sheep monoclonal (1:10.000, GE Healthcare UK, LNA931V/AH); Peroxidase anti-rabbit IgG rat polyclonal (1:10.000, Jackson Immuno-research, 312-006-045).

### Plasmids

All the plasmids used were sequenced by the plasmidsaurus company. The ORP3 plasmid was a gift from the Olkkonen team, and corresponds to the full-length human ORP3 isoform 1a 887 amino acids (Lehto et al., 2005). We produced a collection of ORP3 mutants by editing this plasmid. All ORP3 plasmid exhibit kanamycin resistance and harbor either enhanced GFP (GFP) or mCherry N-terminal tags. The GFP-ORP3 Δ-PH mutant was generated using a synthetic gene, with a deletion of aa 51-145 aa of ORP3 corresponding to the PH domain, and cloned using HindIII/ Bbsi insertion sites. The GFP-ORP3 FFAT^m^ plasmid, with mutations in the FFAT motif (EFFDAQE → EVVAAQE) was generated by PCR mutagenesis, and inserted using EcoNI/XagI restrictions enzymes. The GFP-ORP3 FFAT-like^m^ plasmid, with mutations in the FFAT-like motif of ORP3 (HFFSGST → HVVAGST) was generated by PCR and inserted in the ORP3 plasmid with HindIII/BbsI restriction enzymes. The GFP-ORP3 FFAT-like^m^ and FFAT^m^ double mutant plasmid were generated by PCR and inserted using BbsI and Kpn1 enzymes. Likewise, the GFP-ORP3 phospho-mimetic FFAT-like and GFP-ORP3 ΔORD mutants were generated by PCR. The GFP-ORP3 ΔDLSK mutant was generated by insertion of a synthetic sequence of the ORD domain deleted for the gacctgtccaag (DLSK) sequence, and cloned using BbsI/AflII enzymatic restriction.

PCR mutagenesis was performed using the following 5’ to 3’ templates (capital letters indicate mutation sites). p-ORP3-FFAT^m^ (5’ gactccctttctgagGtGGttgCtgctcaggaagttctgttatctccaag 3’), p-ORP3-FFAT-like^m^ (5’ tccacatgaagttaaccacGtGGtGGcCgggtccaccatcacagactcttcatc 3’), p-ORP3-FFAT-like-phospho-mimetic (5’ catgaagttaaccactttttcGACggAtccGATatcacagact -cttcatctgg 3’), p-ORP3-Δ-ORD ( 5’ catctgaGAATTCAACATCCTGAGGAACA -ACATCGG 3’). The mCherry-VAPA and mCherry-VAPA^KD/MD^ plasmids were generous gifts from Fabien Alpy (IGBMC, Illkirch). The GFP-VAPB and GFP-P4M-SidM probe plasmids were obtained from Addgene (#108126 and # 514711 respectively). GFP-PH-PLCδ was a kind gift from Francesca Giordano (I2BC, Gif-sur-Yvette, France). The GFP-MAPPER probe was a gift from Jen Liou (UT Southwest Medical Center, Dallas), also available as Addgene plasmid #117721. RFP-KDEL was a gift from Nihal Altan-Bonnet (NHLBI, Bethesda).

Silencing ORP3 by RNA interference (siRNA)

The siRNA sequences target the endogenous mRNA 5’ end, upstream of the ORP3 PH domain, absent from the synthetic plasmid. The absence of off-targets was verified using BLAST.

ORP3 siRNA 1 5’ - ccuugauaguggucgggaatt -3’

mock ORP3 siRNA 1 5’ - gugcagauugacggcuagutt -3’

ORP3 siRNA 2 5’ - gcuuucuaaugaaaguagatt -3’

mock ORP3 siRNA 2 5’ - agaacuauugaucguaaugtt -3’

Note: The ORP3 siRNA 1 and mock ORP3 siRNA were previously used in Letho *et al., 2005*.

Cell line editing with CRISPR/Cas9

The HeLa AP CRISPR-Cas9 ORP3-KO cell line was generated using a single nickase GFP ORP-3 CRISPR/Cas9 KO Plasmid (Santa Cruz), containing a pool of the three following guide RNA (gRNA):

gRNA 1 5’-gggggagatgaattacaccc -3’

gRNA 2 5’-tacctgtcgacttccttgct -3’

gRNA 3 5’-atgcatagaccttgacaccg -3’

These gRNA are targeting sequences in the mRNA sequence of ORP3 prior to the start codon of the wild-type isoform. The control plasmids contain scrambled guide RNA, with no detectable off-target in humans using BLAST. HeLa cells were transfected with either the ORP3 CRISPR/Cas9 plasmid, or the control plasmid using Lipofectamine 2000 (Thermofisher). Due to the absence of antibiotic resistance in the plasmid, positive clonal cell lines were isolated by fluorescence activated cell sorting (FACS) using the GFP signal. The clones were further screened by Western blotting for the absence of ORP3. A secondary screen was performed to select GFP-negative cells (for further use of GFP-tagged proteins). The ORP3-KO stable cell lines were used to generate stable rescue cell lines expressing wild-type GFP-ORP3 or mutants (cf “Plasmids” section). Stable clones were sorted by FACS and verified by Western blotting and fluorescence microscopy. VAPA-KO cell lines were generated by CRISPR/Cas9 using double nickase plasmids (Santa Cruz, sc-424440-NIC) and controls (Santa Cruz, sc-437281), as previously described (Siegfried et al., 2024). The mCherry-PH-PLCδ and mCherry-P4MSidM stable cell lines were generated in three different backgrounds: HeLa AP wild type cell lines, mock ORP3-KO cell lines and ORP3-KO cell lines and sorted to generate clonal populations using FACS. The cell lines were verified by Western blotting and microscopy.

### Cell synchronization

The protocol used for prometaphase synchronization was described in (Zieve et al., 1980). The cells were seeded to reach 80% confluence on the first day of treatment. Cells were treated with thymidine 2mM (Merck KGaA) for 24h, to ensure blocking in S-phase. The thymidine block was released for 3h in Phosphate Buffered Saline (PBS, Gibco) and washed to allow cell-cycle progression to prophase. 100 ng/ml Nocodazole (Merck KGaA) was added to the culture medium for 12h to ensure the mitotic block in prometaphase. The block was then released by PBS wash, allowing cell progression through mitosis.

### Western blotting

Cells were grown to confluence in 6-well plastic dishes (TPP) and washed 3 times with PBS. Cells were lysed for 10 min at 4°C in 200µl lysis buffer 50 mM Tris pH 8.5 (CarloErba SDS, 1560517), 150 mM NaCl (MERCK, 1.06404.1000), 1X protease inhibitor cocktail with EDTA (Roche, 05056489001), 0.5% triton-X100 (Sigma, T9284-100ML). Lysates were centrifuged at 13000 g for 20 min at 4°C and supernatants were collected without the insoluble fraction. Protein concentrations were quantified in 96-well plates (TPP) using a Spectramax imager, using the Bradford assay (BioRad). 20µg total protein was diluted in loading buffer (Fermentas, R0891) and denaturated at 95°C for 10 min. The PageRuler Plus Prestained Protein Ladder (Thermoscientific, 26619) was used as a size marker. Electrophoresis was performed at 200V in 4-12% Mini-PROTEAN TGX Stain-Free Precast Gels (Bio-Rad) in running buffer containing 25mM Tris, 192 mM glycine (MERCK, 10421025), 0.1% SDS (Euromedex, Eu0660) and transferred at a current of 400mA onto PVDF membranes (Amersham Hybond, GE10600023) in transfer buffer containing 25 MM Tris, 192 mM glycine, 20% ethanol (VWR, 20824.365). Membrane blocking was performed using blocking buffer with 5% skimmed dry milk (Régilait) in PBS containing 0.1% Tween-20 (Biosolve, 20452335). Primary antibodies were incubated incubated overnight at 4°C. After three washes with blocking buffer, secondary HRP antibodies were diluted in blocking buffer and incubated for 1 hour, then washed three times. Antibodies were detected using the Supersignal west femto maximum sensitivity substrate (Amersham, KB011) and visualized with a ChemiDoc chemiluminescence detection system (BioRad). Quantification of Western blots was performed by densitometry by comparing the band of interest with a housekeeping gene using the Gel analyzer plugin of Fiji imaging software (version 2.16.0).

### Immunoprecipitation

HeLa cells were seeded the day before the experiment, then lysed (cf “Western blotting" section). The cell lysates were incubated overnight with IgG Sepharose beads (GE Healthcare, 17-0969-01) coated with anti-VAPA primary antibodies (cf “Antibodies” section). After 5 washes with lysate buffer, bound proteins were denatured for 10 min at 95°C, and subsequently loaded for Western blotting.

### CIP dephosphorylation

HeLa cells were seeded the day before the experiment and synchronized (cf "Cell Synchronization" section). After lysis, the proteins supernatant was treated with CIP phosophatase for 1 hour at 37°C.

### Live-cell imaging

For live-cell imaging cells were grown on segmented IBIDI µ-Dish 35 mm (IBIDI), to ensure simultaneous and identical imaging conditions for the different experimental conditions. 90 min prior to imaging, sample media were changed for FluoroBrite™ DMEM, supplemented with 10% fetal bovine serum and 100 units/ml of penicillin. When needed, this was completed with fluorescent DNA, tubulin or actin probes (Spirochorme, cf “Probes for live-cell imaging” section). Microscopy was performed in a closed chamber set to 37°C under 5% CO_2_ atmosphere on a Yokogawa CSU-X1 Spinning Disk confocal mounted on an inverted motorized Axio Observer Z1 (Zeiss) and equipped with a sCMOS PRIME 95 (photometrics) camera and a 63X oil-immersion objective (1.4 NA). Z-stack images were acquired at a 2µm z-steps, except for tubulin when z-steps were 0.5 μm. Images were acquired using the MetaMorph software (Version 7.10.5.476). To limit phototoxicity, movies were acquired for 2 hours with a time resolution of 3 min. Image processing and analysis were done in Fiji imaging software.

### Immunofluorescence microscopy

Cells were seeded on 18mm diameter glass coverslips (#1.5, VWR, 631-0153) at 60% confluence. The next day, cells were washed with PBS. For formaldehyde fixation (Thermo scientific, 28908), cells were fixed in a 4% formaldehyde diluted in PBS for 15 min at room temperature. For methanol fixation (VWR, 20847.295), cells were fixed using pre-chilled methanol (-20°C) for 15 min. Cells were then washed 3 times with NH4Cl 50mM (VWR, 21236.291) in PBS, then incubated in permeabilizing solution containing 0,5% Bovine Serum Albumin (BSA, Sigma, A3059-100G) and 0,1% Triton (Sigma, T9284-100ML) in PBS for 15 min. Cells were incubated with primary antibodies diluted in the incubation solution for 1 hour at room temperature. Samples were washed 3 times using the permeabilizing solution. Cells were next incubated with the secondary antibodies for 1 hour at room temperature used in the permeabilizing solution with cross-adsorbed Alexa 488 and/or Alexa 555 anti-mouse/anti-rabbit IgG secondary antibodies (cf antibody section) and washed in PBS. Cells were stained with DAPI (Sigma Aldrich, ref: D9542) for 5 min in PBS, washed In PBS and mounted on slides using 8µl VECTASHIELD® Antifade Mounting Medium (Vector laboratories), and sealed using nail polish. Slides were images on a Yokogawa CSU-X1 Spinning Disk microscope (cf fluorescent microscopy section).

### Electron microscopy

In order to examine the distribution of membrane contact sites between the ER and the plasma membrane, Serial block-face Scanning electron microscopy (SBF-SEM) EM was used. This technique allows 3D imaging of cells and tissues at the ultrastructural level. HeLa Cells were fixed in 3% formaldehyde (Thermo scientific, 28908), 1% glutaraldehyde (Electron microscopy sciences, 16400) for 1 hour at room temperature and prepared for SBF-SEM as previously described (Lachat et al., 2022). For imaging, a TeneoVS SEM (ThermoFisher Scientific) was used with a beam energy of 2.7 kV, 100 pA currents, in LowVac mode at 40 Pa, a dwell time of 1 μs per pixel at 10 nm pixel size. Sections of 100 nm were serially cut between images.

### Microscopy Image analysis

The percentage of multinucleated cells within a population was assessed by imaging formaldehyde-fixed samples on coverslips, stained with DAPI for 5 min before fixation. For each experimental condition, 20 regions of the coverslip were randomly acquired. The ratio of multinucleated/mono-nucleated cells was calculated using the Fiji cell counter plugging. Only cells fully visible in the 3D-field of view with non-ambiguous nuclei in the different z-stacks were analyzed. Division aberrations and cell division timing were characterized by live imaging of dividing cells stained with SiR-DNA (λ674) probe. Cells were incubated 1 hour before imaging in FluoroBrite™ DMEM (Gibco) supplemented with 10% fetal bovine serum (Gibco), 100 units/ml of penicillin, SiR-DNA (λ674) and Spy-Actin(λ505). PI(4,5)P2 quantification in mitotic cells, was performed by live cell imaging of mCherry-PH-PLCδ stable cell lines. The intensity was measured along the cell contour starting at one pole of the dividing cell. For quantification purposes (averaging between cells), the total length of the cell contour was normalized to 100. The same approach was used to measure actin intensity at the cortex in late mitosis. Colocalization between ORP3 and VAPA, or ORP3 and KDEL was assessed by using the Pearson colocalization module of the Fiji imaging software. Before quantification, images were filtered (Median filter r=1, Gaussian blur r=1). To restrict the localization analysis to the cytosol where both proteins are present, a mask corresponding to ORP3 distribution was created, and added to the colocalization settings. Furrow size was measured on 15 z-stack slides, where z-resolution was set at 0.5μm.

### Graphs and statistical analysis

Quantifications were performed using two-tailed Student’s t-test assuming equal variances to compare the distributions. * p < 0.05; ** p < 0.01. *** p < 10^-3^. The graphs were generated using templates available on bio render.

